# Prestimulus alpha power is related to the strength of stimulus representation

**DOI:** 10.1101/2020.02.04.933382

**Authors:** Louise C. Barne, Floris P. de Lange, André M. Cravo

**Affiliations:** Centro de Matemática, Computação e Cognição, Universidade Federal do ABC (UFABC), São Bernardo do Campo, SP, Brasil; Donders Institute for Brain, Cognition and Behaviour, Radboud University, The Netherlands

**Author notes:** **Corresponding author** André M. Cravo, Centro de Matemática Computação e Cognição, Universidade Federal do ABC, Rua Arcturus, 03. Bairro Jardim Antares. São Bernardo do Campo - SP - Brasil. CEP 09606-070. **Author contributions statement:** L.C.B., F.P. L. and A.M.C. conceived the experiment. L.C.B. performed the experiments and analysed the data. All the authors wrote and reviewed the manuscript.

**Keywords:** spatial attention, alpha oscillations, multivariate pattern analysis, EEG

## Abstract

Spatial attention can modulate behavioural performance and is associated with several electrophysiological markers. In this study, we used multivariate pattern analysis in electrophysiology data to investigate the effects of covert spatial attention on the quality of stimulus processing and underlying mechanisms. Our results show that covert spatial attention led to (i) an anticipatory alpha power desynchronization; (ii) enhanced stimuli identity information. Moreover, we found that alpha power fluctuations in anticipation of the relevant stimuli boosted and prolonged the coding of stimulus identity.

## Introduction

Attention is essential to select and process sensory stimuli in the environment. One of the main difficulties in studying attentional selection in humans is assessing the quality of processing of attended and ignored stimuli. Measures such as the amplitude of electrophysiological evoked responses (i.e. N1, P1) have been used as a reflection of this processing. However, a recent study has questioned this view since the amplitude of the evoked response was not correlated with a boost in the target representation.^1^. Moreover, how well the unattended stimulus is processed is particularly hard to assess, given that participants are typically asked not to respond to these events.

One possibility is to use methods which evaluate neural processing without necessarily requiring a behavioural response. Recent studies have relied on multivariate pattern analysis (MVPA) techniques to analyse EEG/MEG responses.^2–4^. In general, these methods are based on the theoretical basis that the MEG/EEG signal reflects coupled dipoles activity in the underlying neural circuitry. Although the specific dipole activity may not be identified, their summation would result in different patterns of activity across MEG/EEG sensors^2–4^. These MVPA methods have been increasingly used to understand the effects of attention and its underlying mechanisms. Several studies have shown that spatial attention modulates stimulus representation in working memory^5–7^ and sensory information input^8^. Moreover, temporal attention seems to enhance relevant sensory information input, creating a temporal protection window against distractors^1^, and boosting its representational content.

Another pre-activating mechanism commonly described in the literature of spatial attention is the modulations in the spectral characteristics of the alpha (8-16 Hz) band^9^. This oscillatory activity is associated with cognitive functions such as visual attention^10^, working memory^11^ and cognitive load^12^. For example, anticipatory lateralized desynchronisation of alpha power is correlated with spatial expectations of relevant upcoming stimuli in a particular hemifield^, 10, 13^. Studies suggest that alpha oscillations can play a role in information processing by suppressing the activity of particular neural populations involved with the irrelevant stimuli feature/spatial location processing, and by selecting the inhibition release of task-related neural populations^14, 15^. These anticipatory states reflected by the prestimulus power of low-frequency oscillations are also related to modulations in the amplitude of sensory event-related potentials^16, 17^. The general relationship between oscillatory activity and evoked responses is still not clear, and different explanations have been proposed on how these two relate^17, 18^. In the specific case of anticipatory alpha power and evoked responses, there is evidence for both positive and negative correlations. Moreover, the mechanisms by which alpha oscillations modulate perception is also unknown. Recent studies have challenged the view that alpha power modulates perceptual precision and have suggested that it increases perceptual^19^ or decision bias^20, 21^.

In the present work, we combined frequency and multivariate pattern analyses to investigate: (1) how covert spatial attention modulates stimulus representation; (2) whether and how different EEG markers are modulated. We adapted previous tasks that presented targets of different modalities rhythmically^22, 23^ to investigate whether spatial attention induces generic anticipatory mechanisms. Critically, we combined both analyses to investigate how anticipatory mechanisms of spatial attention can influence stimulus processing. We found that anticipatory alpha can modulate performance by modulating baseline excitability and improving the precision of stimulus representation.

## Materials & Methods

### Data and script availability

Data (raw and analysed), presentation and analysis scripts are publicly available at (https://osf.io/wbhc8/). The procedures and analysis were not pre-registered prior to the research being conducted

### Participants

Twenty participants (age range, 18-27 years; 11 female) gave informed consent to participate in this study. The sample size was based on previous EEG studies^6, 8, 24^. Data from one participant was excluded from the final analysis due to excessive noise in the EEG signal. All participants had normal or corrected-to-normal vision and were free from psychological or neurological diseases (self-reported). The experimental protocol was approved by the Research Ethics Committee of the Federal University of ABC.

### Apparatus

Stimuli were presented using the Psychtoolbox v.3.0 package for MATLAB on a 20-inch CRT monitor with a vertical refresh rate of 60 Hz, placed 50 cm in front of the participant. EEG was recorded continuously from 64 ActiCap Electrodes (Brain Products) at 1000 Hz by a QuickAmp amplifier (BrainProducts). All sites were referenced to FCz and grounded to AFz. The electrodes were positioned according to the International 10-10 system. Additional bipolar electrodes registered the electrooculogram (EOG).

### Procedure

Each block began with the presentation of a central fixation point (white, 0.4°) and two circles (horizontal eccentricity of 5°, size of 3°), one at the left and the other at the right side of the fixation point (Figure 1A). Each experimental block consisted of two streams of stimuli, presented inside the circles in one of the visual hemifields. Stimuli presentation was one at a time, alternating between visual hemifields, always starting at the left side, with an SOA of 500 ms between them (2Hz frequency). Participants had to identify the oddball stimuli within a stream of standard stimuli presented at a specific side by pressing a button with their right index finger. Standard visual stimuli were vertical/horizontal oriented Gabor patches (3 degrees of visual angle) with a spatial frequency of 2 cycles per degree (cpd). The oddball stimuli (target) were vertical/horizontal oriented Gabor patches (3°) with a higher spatial frequency defined by an adaptive procedure. There were 70% of standards (half vertical and half horizontal) and 30% of targets. All stimuli lasted 0.1 s.

**Figure 1.**
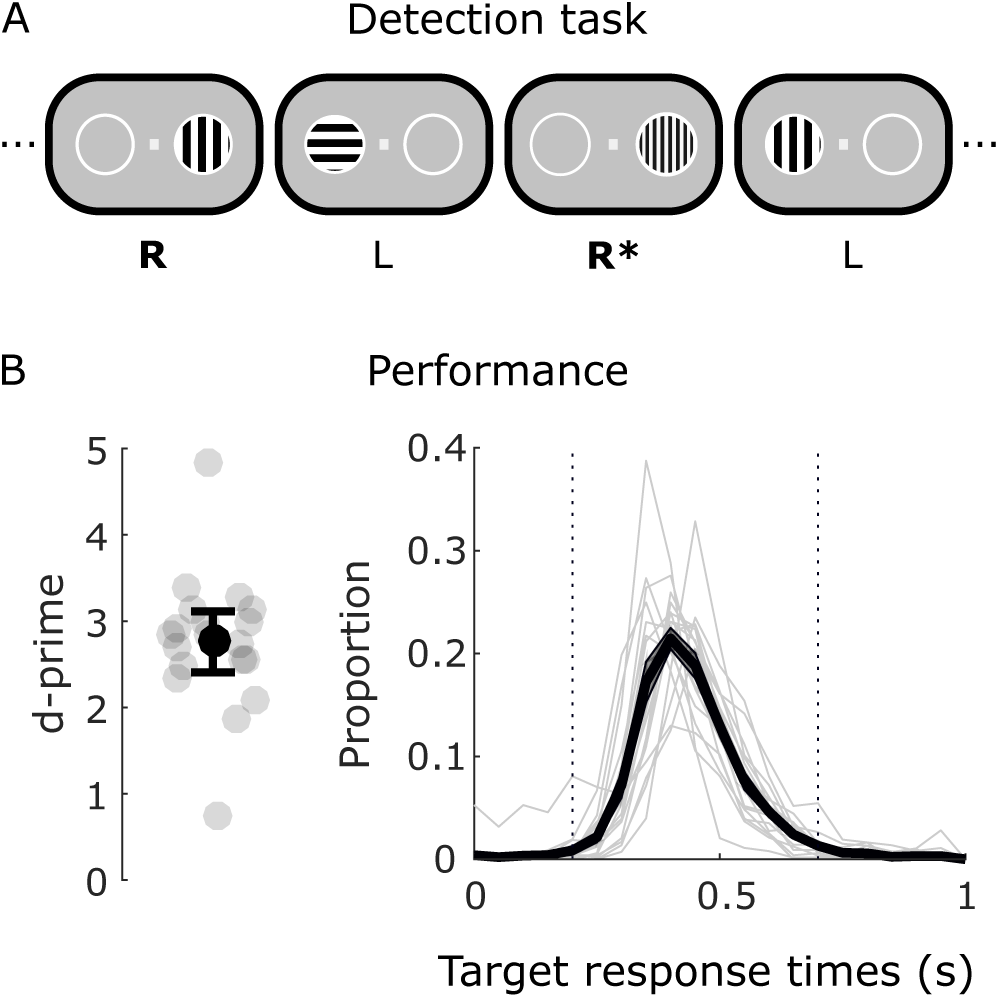
A) Each experimental block began with the presentation of a fixation point. Visual stimuli appeared in sequence, in alternated sides, separated by a stimulus onset interval of 0.5 s. In different blocks, participants had to respond to the oddball target either on the right or left stream. The oddball stimuli (target) were Gabor patches with a higher spatial frequency defined by a staircase procedure. There were 70% of standards and 30% of targets. All stimuli lasted 0.1 s. B) Left: Distribution of participants’ d-primes (circles) and its respective errorbar (mean, S.E.M). Right: Reaction times for targets (mean, S.E.M). Dashed lines represent the interval that was considered as hits. Lighter lines represent individual response times.

Each block was preceded with an instruction screen indicating in which visual hemifield (right or left) the participant had to detect targets, thus resulting in two main experimental situations: attend to the right or left hemifield. Volunteers were instructed to fixate at the fixation point, but also to allocate their covert attention to the indicated side. Each block consisted of 120 stimuli, 60 for each location, with 18 targets. Therefore, for each location, there were 21 vertical and 21 horizontal oriented standard and 9 horizontal and 9 vertical oriented oddball patches. The order was pseudo-randomized considering no two consecutive oddball stimuli. There were 36 blocks (18 blocks for each condition). Block types were presented in a pseudorandom order (not more than three same attended side blocks in a row).

The session began with volunteers performing a training session, which consisted of one block of the experimental procedure followed by a weighted up-down staircase procedure^25^. The staircase block was shorter, with 80 stimuli. After every 12 attended side stimuli presentation, the spatial frequency value was recalculated based on accuracy. A psychometric function was fitted to the data to obtain the spatial frequency corresponding to 75% of correct responses.

Interleaved with the experimental blocks, participants performed a localizer task. The task was to detect a colour change in the fixation point (from white to yellow) while the standard stimuli were presented sequentially (SOA of 500 ms) and randomly at left and right sides. Five blocks were presented after every nine main experimental blocks, totalling 15 localizer blocks and 900 standards (50% vertically oriented) presented at each side in total. The localizer was initially designed to be used as a training set to the classification procedure in the main experiment. However, this procedure did not work well, and scores were not above chance within the localizer blocks. For this reason, we have decided not to include this analysis in the paper. Data, analysis and a brief discussion of the localizer data can be seen at https://osf.io/ca2fd/.

### EEG pre-processing

EEG pre-processing was carried out using BrainVision Analyzer (Brain Products). All data were down-sampled to 250 Hz and re-referenced to ear electrodes. For eye movement artifact rejection, an independent component analysis (ICA) was performed on filtered (Butterworth Zero Phase Filter between 1 Hz to 30 Hz) data. Eye related components were identified by comparing individual ICA components with EOG channels and by visual inspection.

### EEG analysis

EEG analyses were based only on standard stimuli to avoid contamination from motor responses. Standard stimuli presented 500 ms before or 200 ms after a motor action were also excluded from further analyses. Stimuli were separated, in different analyses, based on Attention (attended/ignored), stimuli hemifield occurrence (right/left), and stimuli orientation (vertical/horizontal).

ERP consisted of a filtered (Butterworth Zero Phase Filter between 0.05 Hz to 30 Hz) segmented (−0.25 to 0.495 s, where 0 is the onset of the stimulus) and baselined (−0.25 to 0 s) data. For the frequency analysis, no-filters were used and data were segmented between −1.95 and 1.95 s relative to stimulus onset. Time-frequency analysis was based on a short-time Fourier transform of Hanning tapered data. We estimated frequencies between 4 and 40 Hz, using a frequency dependent time window of 5 cycles that was advanced over the data in 20-ms steps. Power was logarithmically transformed in the end. Analyses were performed using FieldTrip toolbox^26^ (http://fieldtriptoolbox.org).

For the MVPA analysis, ERP data from all conditions and all electrodes were exported from MATLAB to Python 3 Jupyter Notebook (5.0.0) to decode stimulus orientation using Scikit-learn library. Classification of grating orientation (vertical or horizontal) was performed separately for each attention condition and side (attended stimulus presented at the left hemifield, ignored stimulus at the left hemifield, attended stimulus at the right hemifield, ignored stimulus at the right hemifield). All EEG channels were used as features for the classification procedure. The classification pipeline consisted of a leave-one-block-out cross-validation (18 blocks) procedure. Training and test data were normalised between 0 and 1 based on the training data, and a logistic classifier with a L2 regularisation was used with all parameters set to the default in Scikit. For each participant, time point and model, the logistic regression pipeline returned labels’ probabilities of the trials and the area under the receiver operating characteristic curve (AUC) was calculated as performance score. Afterwards, left and right scores were averaged with respect to attended and ignored conditions.

Critically, the experiment was designed so that the: i) feature that defined whether the stimulus was a target or not (spatial frequency) was orthogonal to the feature to be decoded (orientation); ii) both features (spatial frequency and orientation) shared a common a similar neural mechanism, meaning that participants would not be able to ignore the feature to be decoded while trying to perform the task.

## Statistics

### Relation between reaction times and pre-stimulus alpha power

We ran two generalised linear mixed-effects models in MATLAB (glme), a linear model using log-transformed reaction times as the dependent variable and a second binomial one considering hits as the dependent variable. Pre-stimulus alpha power was the fixed factor, and participants’ slope and intercept in alpha power were the random factors.

### EEG

Statistics of one-dimensional classification scores were performed non-parametrically^27^ with sign-permutation tests. For each time-point, the decoding value of each participant was randomly multiplied by 1 or −1. The resulting distribution was used to calculate the p-value of the null-hypothesis that the mean discrimination-value was equal to 0.5. Cluster-based permutation tests were then used to correct for multiple comparisons across time using 1000 permutations, with a cluster-forming threshold of p <0.05. The significance threshold was set at p <0.05 and all tests were two-sided. Similar results were obtained when performing univariate t-tests and using an FDR based correction on the estimated p-values. These statistical analyses were performed for the ERP, time-frequency and Fourier Transform analysis in MATLAB. Moreover, cluster-based analysis implemented in the MNE library (version 0.18.2, default parameters) was performed to calculate whether decoding scores were statistically different from the null-hypothesis (AUC = 0.5), and to calculate whether attended and ignored decoding scores were more likely drawn from different probability distributions (function: permutation cluster 1samp test).

## Results

### Behavioural results

Participants’ performance were computed considering hits as responses between 200 ms to 700 ms after a target was presented at the attended side (Figure 1B, right). All other responses were considered false alarms. Participants were able to perform the detection task (Hit Rate: mean = 0.75, SEM = 0.033, min = 0.344, max = 0.976; and d’: mean = 2.759, SEM = 0.181, min = 0.727, max = 4.84, Figure 1B, left). Additionally, there was no significant effect of side on d-prime (mean ± SEM, left: 2.824 ± 0.219; right: 2.733 ± 0.167; t(18) = 1.023, p = 0.32).

### Event related potentials

To investigate the effects of attention on event-related potentials, we focused on contralateral electrodes relative to the visual hemifield of the standards stimuli. Specifically, we focused on parieto-occipital electrodes (Right hemifield conditions: PO9/PO7/ PO3/O1; Left hemifield conditions: PO10/PO8/PO4/O2), baselined between −50 to 50 ms. Although attended stimuli evoked a stronger negative peak around 180 ms (Figure 2, left), there was no significant difference between attended and ignored conditions (highest negative cluster-stat = 51.3833, p = 0.18, time = 0.172 to 0.2, highest positive cluster-stat = 51.3833, p = 0.068, time = 0.42 to 0.5). Although we did not find significant differences between conditions, it is important to notice it is hard to disentangle attended and ignored ERP due to the fixed SOA design and possible component overlapping. However, there was clear activity lateralization due to attention (Figure 2, right).

**Figure 2.**
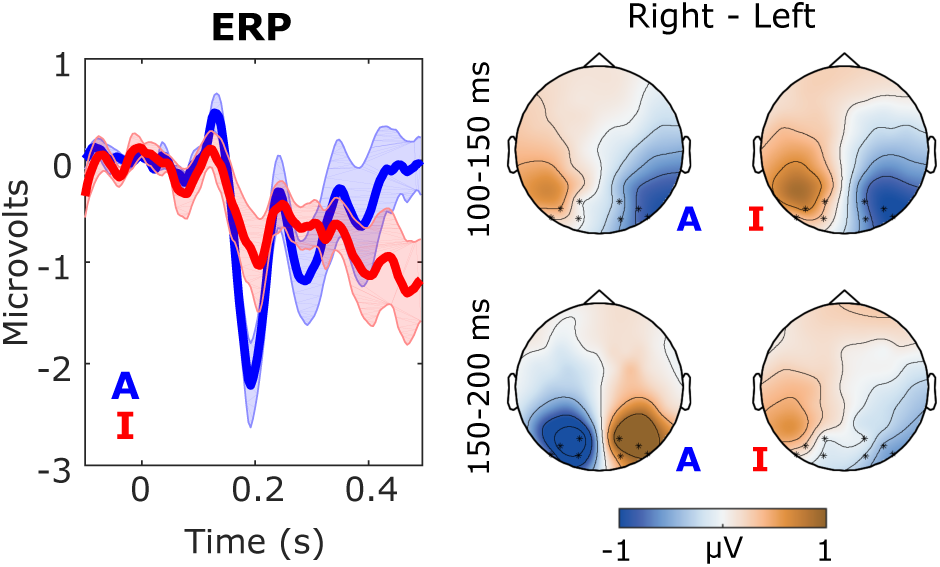
Event related potentials. Left: Activity evoked by attended (blue) and ignored (red) standards. Time point zero corresponds to the onset of stimuli. Right: Topographic plots showing the difference between the average activity of the standard stimuli presented at right and left hemifields when attended and ignored during P1 and N1 time windows (100-150ms and 150-200 ms). Asterisks represent used parieto-occiptal channels.

### Alpha-band oscillations

Alpha band desynchronization is a key marker of attention. Therefore, we compared the power spectrum of attended and ignored conditions. Similar to the ERP analysis, we focused on the same parieto-occipital contralateral electrodes to stimulus presentation when that stream was to be attended or ignored(Figure 3 A). There was a difference in power evoked by attended and ignored stimuli (positive cluster-stats = 1203.6, 1269.2, 1791.6, all p <0.01; negative cluster-stats = 774.26, p<0.004; stats = 502.33, p = 0.027). The clusters’ time period coincides with stimuli occurrence, which is in line with previous work showing that attention modulates the post-stimulus induced responses as the power of low-frequency bands (alpha/beta)^28, 29^. Additionally, there was a significant alpha (9 - 15 Hz) power attenuation when attended standard stimuli were about to be presented compared to ignored ones.

**Figure 3.**
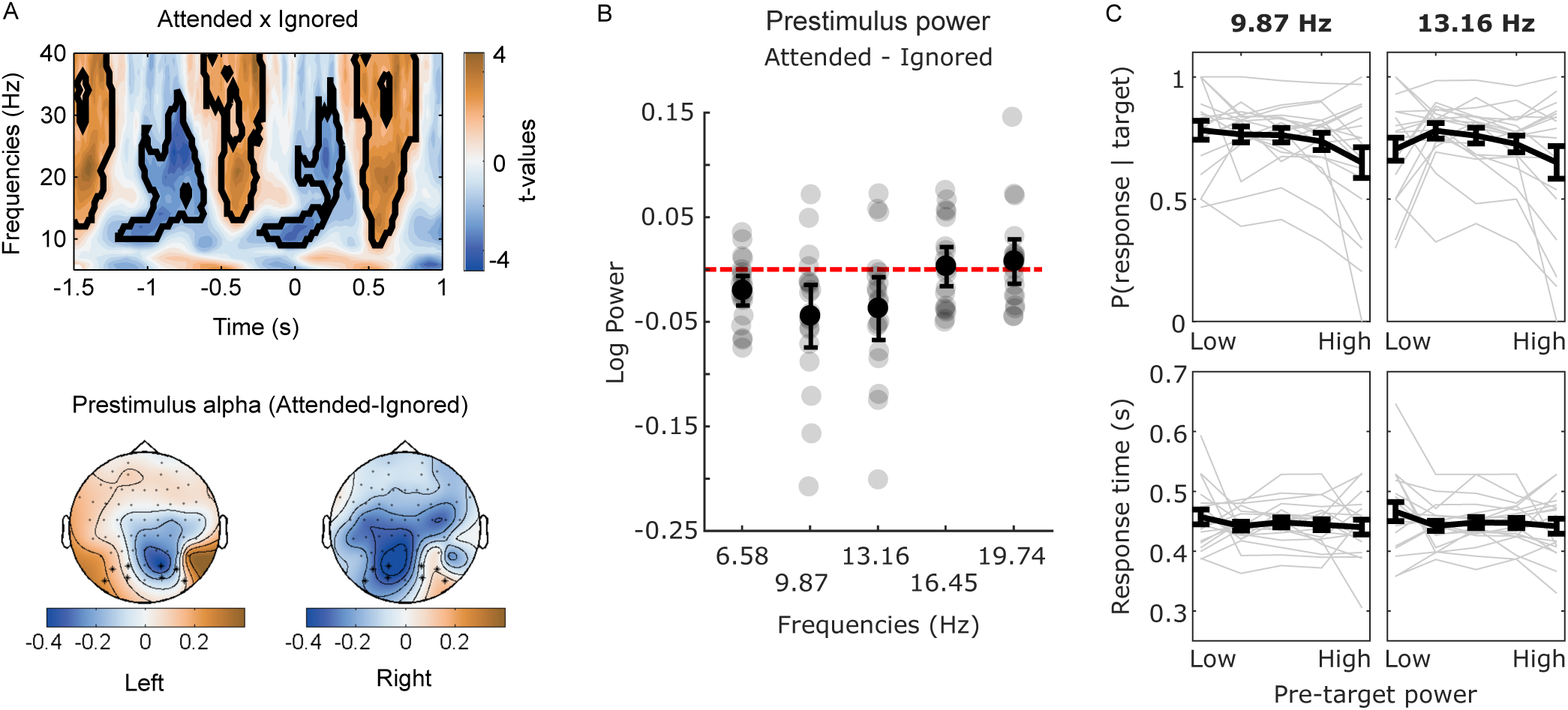
A) Top: Power spectrum paired t-test comparison between attended and ignored stimuli. Black contours indicate significant clusters. Bottom: Alpha power (dB) topography from the difference between attended and ignored stimuli at a prestimuli time period (−0.5 to 0 s). Asterisks represent used parieto-occiptal channels. B) Error bars (mean, S.E.M) showing the difference in prestimulus power between attended and ignored conditions at frequencies around the previous significant band. Individual differences are represented as dots in the background. B) Error bars (mean, S.E.M) illustrating hits and reaction times for the pre-target alpha power bins. For graphical purposes, alpha power was binned in five bins (with a similar number of trials) based on prestimulus alpha power and the behavioural output was calculated for each bin. Individual responses for different bins are represented in the background.

In a second step, we restricted our analysis on prestimulus anticipatory alpha power. We ran a Fast Fourier Transform with a Hanning taper on a shorter epoch (−0.3 to 0 s relative to stimulus onset). We targeted frequencies around the previous significant band. Given the window duration and frequency resolution (from 2 to 6 cycles), we were able to compare the prestimulus power for attended and ignored stimuli of five frequencies: 6.58, 9.87, 13.16, 16.45 and 19.74 Hz. Alpha band frequency of 9.87 Hz showed a significant anticipatory attentional effect (Figure 3 B, 6.58 Hz: attended = 0.473 ± 0.049, ignored = 0.494 ± 0.045, t(18) = −2.827, d = −0.649, p = 0.055; 9.87 Hz: attended = 0.550 ± 0.069, ignored = 0.595 ± 0.069, t(18) = −2.922, d = −0.670, p = 0.045; 13.16 Hz: attended = 0.356 ± 0.073, ignored = 0.394 ± 0.073, t(18) = −2.449, d = −0.562, p = 0.125; 16.45 Hz: attended = 0.070 ± 0.075, ignored = 0.067 ± 0.073, t(18) = 0.287, d = 0.066, p >0.9; 19.74 Hz: attended = −0.084 ± 0.064, ignored = −0.091 ± 0.064, t(18) = 0.696, d = 0.160, p >0.9, p-values corrected for multiple comparisons using Bonferroni).

To evaluate whether prestimulus alpha power was correlated with performance, we ran two separate mixed-models: one investigating how alpha power modulated reaction times to correct target detection and one investigating how alpha power modulated target detection (hit rate) for 9.87 Hz and for 13.16 Hz (Figure 3 C). We did not find a significant effect of alpha power on reaction time (9.87 Hz: alpha fixed effect, t = 0.404, p = 0.686, 13.16 Hz model: alpha fixed effect, t = −0.179, p = 0.858), nor hit rate (9.87 Hz model: alpha fixed effect t = −1.76, p = 0.078, 13.16 Hz model: t = −1.903, p = 0.057).

### Multivariate pattern analyses

We used an L2-regularised logistic regression to test whether orientation information could be decoded in different conditions. We trained and tested separately for each attention and side and, subsequently, we averaged the scores separately for attended and ignored conditions (Figure 4 A). We found that classification was above chance levels for attended stimuli (Attended right: time = 0.184 to 0.224, cluster-stat = 41.351, p = 0.031; Attended left: time = 0.144 to 0.18 and 0.384 to 0.408, cluster-stats = 41.19 and 26.72, p = 0.006 and p = 0.034; Attended [mean], time: 0.148 to 0.224, 0.26 to 0.324, 0.356 to 0.428, cluster-stats = 81.23, 63.11, and 71.11, p = 0.002, 0.002, 0.002). There were no significant clusters in the Ignored condition. A second cluster-based analysis comparing Attended and Ignored scores was performed and showed that attended stimuli had higher classification scores than ignored stimuli (time: 0.144 to 0.22 s, cluster-stat = 211.332, p = 0.018).

**Figure 4.**
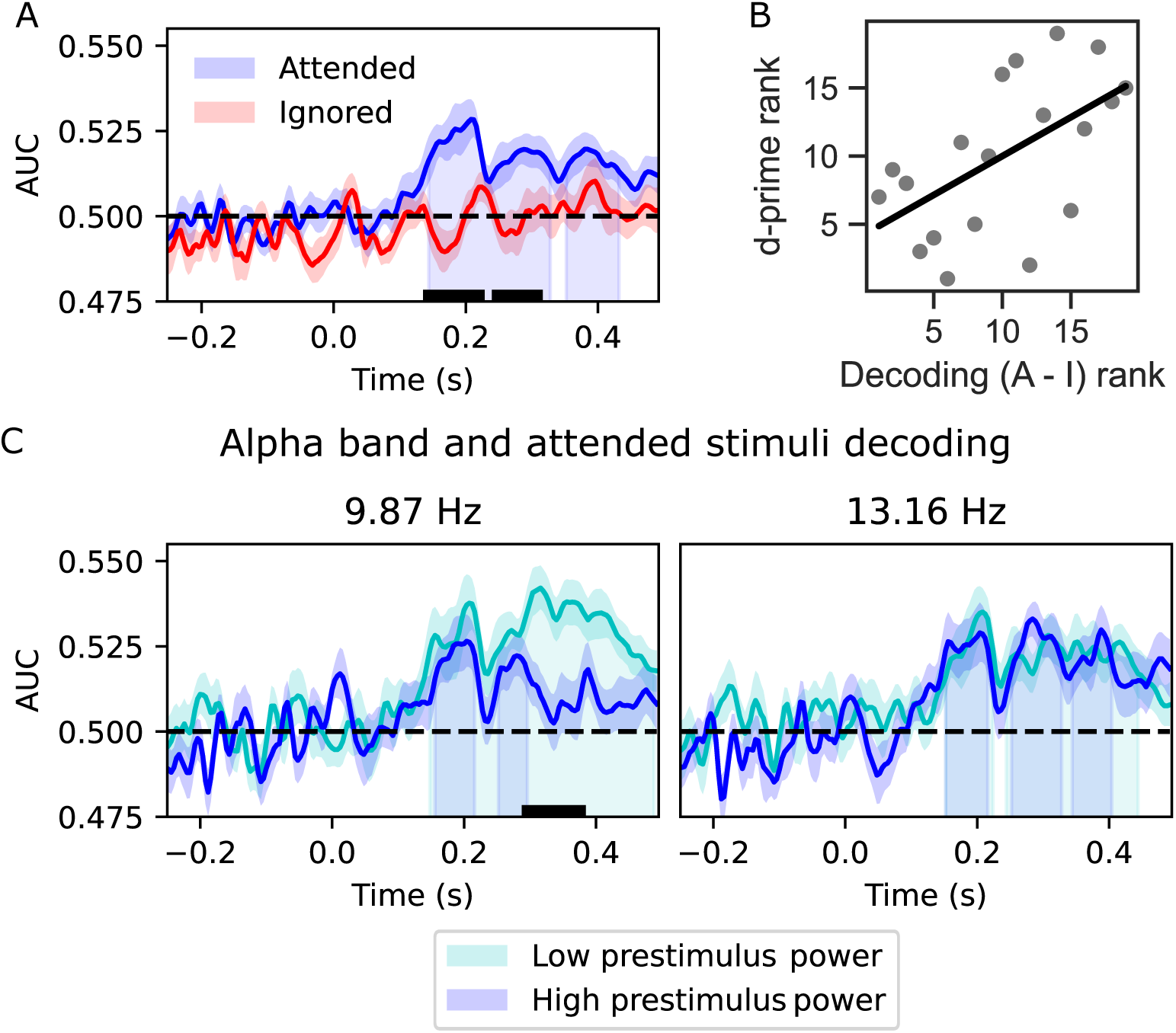
A) Decoding scores across different experimental conditions. Red (ignored) and blue (attended) lines represent mean and error bars represent standard error of the mean. Time zero is the moment in which a standard was presented. Shadows under the curve represent clusters where classification was better than chance. The black line at the left panel shows the cluster period in which classification was higher in attended than in ignored conditions. B) Scatterplot showing the cross-subject correlation between d-prime and the difference in decoding scores between attended and ignored conditions. C) Decoding scores for attended stimuli (mean, S.E.M) grouped into low (cyan) and high (darker blue) prestimuli alpha power. Grouping was performed for both frequencies at alpha band: 9.87 Hz and 13.16 Hz. The black line shows the cluster period in which classification was higher in low alpha power trials than in high alpha power trials, and shadows under the curve represent the statistical clusters different from scores at chance.

To investigate whether higher differences between attended and unattended representations lead to better performances, we computed a across participants (n=19) correlation between d-prime and the difference between attended and ignored decoding scores. We found that attentional manipulation on decoding scores was correlated with participants’ d-prime values (Spearman correlation = 0.57, p = 0.010, Figure 4 B. Similar correlation results were obtained using other methods such as Kendall’s Tau, Sheppard’s Pi and percentage bend correlation).

Given that attention modulated both alpha power (stronger effect at 9.87 Hz) and classification accuracy, we tested whether prestimulus alpha desynchronisation was correlated with an increased classification. To test this hypothesis, we performed a median split on attended stimuli based on their prestimulus alpha power (9.87 Hz and 13.16 Hz) and recalculated the decoding score (AUC) for each participant and time point (Figure 4 C). For the low alpha band (9.87 Hz), a direct comparison between prestimulus power conditions showed a higher decoding score for stimuli preceded by low alpha power (cluster-stat = 63.209, p = 0.008, time = 0.30 to 0.376). Results for the higher frequency of alpha band (13.16 Hz), however, showed no modulation (highest cluster-stat = 10.617, p = 0.495, time = −0.192 to −0.18). Taken together, our results suggest that the frequency in which there was a stronger modulation of attention (9.87 Hz) was also correlated with higher classification scores.

## Discussion

In the present study, we investigated the effects of covert spatial attention on sensory information and related mechanisms. Attention resulted in a pre-stimulus alpha desynchronization on contralateral sensors. Using a multivariate decoding approach, we found that spatial attention enhanced stimulus identity coding at the relevant location. Moreover, we found that that stronger alpha desynchronisation boosted the representation of target identity.

Several studies have shown that visual-spatial attention elicits larger P1 and N1 components for attended than for ignored-stimuli^30^. However, P1 and N1 effects seem to indicate distinct attentional mechanisms: while P1 modulation suggests a suppressing mechanism, it has been suggested that N1 modulation represents a sensory gain mechanism^31^. In this study, we did not find a significant effect of attention on event-related potentials. One crucial difference in our study is that events were presented sequentially in a stream. This type of presentation leads to overlaps of event-related potentials which, in turn, might have made it harder to find effects on early components. Moreover, it has been recently suggested that stronger response amplitudes (i.e. ERP, BOLD response) do not necessarily imply enhancement in information about stimulus identity^32^. A recent study investigating temporal attention found a modulation in N1, but no correlation between N1 amplitude and the decoding scores^1^. Therefore, it seems that an increase in signal-noise-ratio is not necessarily linked to boosting information about stimulus identity.

We observed a stronger contralateral alpha decrease in anticipation of attended targets. Our results are in agreement with several studies that have shown that alpha oscillations are correlated with spatial attention^6, 33–37^. There has been an increasing discussion as to whether alpha oscillations are associated with an increase in perceptual processing or if it modulates performance by biasing perceptual^19^ or decisional mechanisms^20, 21^. In recent studies, it has been suggested that decreased alpha power does not improve perceptual acuity, but biases perception by increasing baseline excitability^19–21^. This proposal is based on findings that show that: (1) decreased alpha power was associated with a higher tendency to report the presence of a stimulus, even when a target is absent; (2) alpha power did not affect performance on discrimination tasks^19–21^. In our experiment, the behavioural task was to detect the presence of a target, marked by an increase in the spatial frequency of the Gabor. Although not significant, there was a tendency of higher reportability rates when alpha power decreased, making our results compatible with the perceptual bias proposal.

Our decoding results showed that spatial attention enhanced target identity coding. Even though the decoded feature (stimulus orientation) and the task-relevant feature (spatial frequency) are independent, they share commons mechanisms and attending to one of them seems to affect the other. Within the attended field, decoding was increased in trials with decreased alpha power. Although this finding appears to be more easily explained with the idea of alpha being associated with perceptual precision, it is worth to point out that the increase in decoding was in a later period. This result is consistent with a previous report that found that lower alpha can reduce the immediate interference by a distractor^1^. Given that participants did not make judgements about the orientation of the target, it is not possible, with our current experiment, to be sure whether an increase in decoding would be translated into an increase in the behavioural discrimination.

We conclude that stimulus identity processing is modulated by covert spatial attention and that anticipatory alpha power is associated with such processing.

## Notes

**Funding:** L.C.B was supported by grants #2016/04258-0, São Paulo Research Foundation (FAPESP) and #2018/08844-7, São Paulo Research Foundation (FAPESP). AMC was supported by grants #2017/25161-8, São Paulo Research Foundation (FAPESP). The funders had no role in study design, data collection and analysis, decision to publish, or preparation of the manuscript.

### Competing Interest Statement

The authors have declared no competing interest.

